# Cell numbers, distribution, shape, and regional variation throughout the murine hippocampal formation from the adult brain Allen Reference Atlas

**DOI:** 10.1101/635201

**Authors:** Sarojini M. Attili, Marcos F.M. Silva, Thuy-vi Nguyen, Giorgio A. Ascoli

**Affiliations:** Center for Neural Informatics, Structures, & Plasticity; Krasnow Institute for Advanced Study, George Mason University, Fairfax, VA (USA)

**Keywords:** Hippocampus, cell counts, mouse, image analysis

## Abstract

Quantifying the distribution of cells in every brain region is fundamental to attaining a comprehensive census of distinct neuronal and glial types. Until recently, estimating neuron numbers involved time-consuming procedures that were practically limited to stereological sampling. Progress in open-source image recognition software, growth in computing power, and unprecedented neuroinformatics developments now offer the potentially paradigm-shifting alternative of comprehensive cell-by-cell analysis in an entire brain region. The Allen Brain Atlas provides free digital access to complete series of raw Nissl-stained histological section images along with regional delineations. Automated cell segmentation of these data enables reliable and reproducible high-throughput quantification of regional variations in cell count, density, size, and shape at whole-system scale. While this strategy is directly applicable to any regions of the mouse brain, we first deploy it here on the closed-loop circuit of the hippocampal formation: the medial and lateral entorhinal cortices; dentate gyrus (DG); areas Cornu Ammonis 3 (CA3), CA2, and CA1; and dorsal and ventral subiculum. Using two independent image processing pipelines and the adult mouse reference atlas, we report the first cellular-level soma segmentation in every sub-region and layer of the left hippocampal formation through the full rostral-caudal extent, except for the (already well characterized) principal layers of CA and DG. The overall numbers (∼600k cells in entorhinal cortex, ∼200k in DG, ∼430k in CA1-3, and ∼290k in subiculum) are corroborated by traditional stereological sampling on a data subset and well match sparse published reports.

## INTRODUCTION

Neuronal and glial numbers are an important attribute in the characterization of distinct functional regions of the nervous system. Cell counts and densities vary considerably between and within brain areas as well as across species and life-span development (Bayer et al. 1982; Herculano-Houzel et al. 2011; Long et al. 1998; Meyer et al. 2010). These numbers are moreover susceptible to pathologies, pharmacological treatment, and genetic alterations (Fitting et al. 2009; Insausti et al. 1998; Malberg et al. 2000; Rajkowska 2000). While the ratio between neurons and glial cells is still actively debated (Bahney and Bartheld 2017; Herculano-Houzel et al. 2013; Sherwood et al. 2006), the numbers of neurons in two inter-connected areas relate to circuit convergence and divergence, which are essential design elements of computational processes. Thus, determining the total number of cells in every brain region is fundamental to attaining a comprehensive census of distinct neuronal and glial types (Insel et al. 2013; Kandel et al. 2013; Kim et al. 2017).

The vast majority of reports of cell counts in the neuroscience literature are based on stereological methods (Grady et al. 2003; Schmitz and Hof 2005; West et al. 1991). In these approaches, the number of cells is accurately measured in a small but unbiased proportion of the volume of interest. The numbers for the whole target region can then be extrapolated under the assumption that the sample be representative. Since stereological neuron counting requires the involvement of a human operator to identify cells in the stained tissue (Schmitdz et al. 2014), the procedure is inherently labor-intensive, time-consuming, and, in particular for cell-sparse regions, impractically inefficient (Boyce and Gundersen 2018). Until recently, the only alternative method that could afford routine comprehensive cellular counting in an entire brain region relied on nuclear identification from homogeneous suspensions (Herculano-Houzel and Lent 2005). While this relatively newer technique distinguishes neurons and glia, it makes it necessary to physically dissect each area of interest and cannot measure geometrical features such as cellular size, shape or spatial distribution within the tissue.

Leaping progress in image recognition and the continuous growth in computing power have substantially altered this status quo (Bhanu and Peng 2000; Peng et al. 2013). Specifically, several algorithms were recently designed to enable high-throughput soma detection and analysis (Hu et al. 2017; Kayasandik and Labate 2016; Luengo-Sanchez et al. 2015; Tapias and Greenamyre 2014; Zhang et al. 2018; Quan et al. 2013). Fully automatic cell segmentation modules are also implemented in freely available mainstream software programs such as ImageJ (Schindelin et al. 2015; Schneider et al. 2012) and CellProfiler (Bray et al. 2015; Lamprecht et al. 2007), allowing robust quantification of soma count, location, and geometry in large-scale applications. In parallel with these computational advances, the public availability of the Allen Brain Reference Atlas (Jones et al. 2009; Lau et al. 2008; Sunkin et al. 2012) provided unprecedented digital access to complete image series of Nissl-stained sections along with regional and laminar delineations.

These new neuroinformatics developments offer a potentially paradigm-shifting alternative to stereological sampling. Taken together, automated cell segmentation and comprehensive online sharing of raw histological datasets provide the opportunity for reliable and reproducible quantification of regional variations in cellular number, size, shape, and spatial distribution at whole-system scale. While this strategy is directly applicable to all regions of the mouse brain, we first deploy it here on the closed-loop of the hippocampal formation, consisting of the six layers of medial and lateral entorhinal cortices and the (three-to-five-layered) dentate gyrus, Ammon’s Horn areas CA3, CA2, and CA1, and dorsal and ventral subiculum.

On the one hand, the initial focus on the hippocampal formation reflects the great interest in this structure by the broad neuroscience community (Hasselmo and Stern 2015; Kandel 2004; Moser et al. 2017). On the other, most unbiased stereology techniques have historically been tested on the hippocampus (Boss et al. 1985; Miki et al. 2005; West et al. 1991), and the relative wealth of published information provides a useful validation benchmark. At the same time, even relatively basic questions on the number and density of cell in the mouse hippocampal formation, such as their medial-lateral, anterior-posterior, and laminar distribution in the entorhinal cortex or in the subiculum, remain surprisingly still open.

Using publicly available software and the full images series of Nissl-stained coronal sections from the standard adult mouse brain Allen Reference Atlas, we present the first complete cell-by-cell soma segmentation in all sub-regions and layers through the full rostral-caudal extent of the left hippocampal formation. We validate this computational counting approach in three ways: by reproducing the analysis with two independent image segmentation pipelines; by confirming the results with traditional stereological estimates on a substantial tissue sample (∼10% of total); and by comparing our findings to available data in the published literature. In addition to comprehensive cell counts, the reported quantitative analysis reveals definitive regional variation of soma geometry and spatial occupancy. With this study, we also release open-source all segmented images, the entire database of raw measurements, and our analysis scripts for further community mining.

## MATERIALS AND METHODS

### Image acquisition

Nissl-stained coronal section images of the adult (8-week old C57Bl/6J male) mouse brain containing any part of the hippocampal formation (sections 64 through 104) were downloaded from the Allen Institute for Brain Science’s Mouse Reference Atlas (RRID:SCR_002978) at the highest available resolution (90 dpi). The hippocampal formation is composed of the hippocampus proper, which consists of the dentate gyrus and Cornu Ammonis areas 1-3 (CA1, CA2, CA3), the medial and lateral entorhinal cortices, and the dorsal and ventral portion of the subiculum. The Allen Brain Atlas further delineates these regions into sub-areas and layers, giving rise to 45 hippocampal structures: strata lacunosum-moleculare, radiatum, pyramidale, and oriens of Cornu ammonis 1 and Cornu ammonis 2 (CA1slm, CA1sr, CA1sp, CA1so, CA2slm, CA2sr, CA2sp, CA2so); strata lacunosum-moleculare, radiatum, lucidum, pyramidale, and oriens of Cornu ammonis 3 (CA3slm, CA3sr, CA3slu, CA3sp, CA3so); the molecular, granular, and polymorphic layers of the dentate gyrus (DG-mo, DG-sg, DG-po); layers 1, 2, 2a, 2b, 2/3, 3, 4, 4/5, 5, 6a, and 6b of the lateral entorhinal cortex (ENTl1, ENTl2, ENTl2a, ENTl2b, ENTl2-3, ENTl3, ENTl4, ENTl4-5, ENTl5, ENTl6a, ENTl6b); layers 1, 2, 2a, 2b, 3, 4, 5, and 6 of the dorsal zone of the medial entorhinal cortex (ENTm1, ENTm2, ENTm2a, ENTm2b, ENTm3, ENTm4, ENTm5, ENTm6); layers 1, 2, 3, and 5/6 of the ventral zone of the medial entorhinal cortex (ENTmv1, ENTmv2, ENTmv3, ENTmv5-6); and strata moleculare, pyramidale, and radiatum of both the dorsal and ventral parts of the subiculum (SUBd-m, SUBd-sp, SUBd-sr, SUBv-m, SUBv-sp, SUBv-sr). We omitted the pyramidal layers of Cornu ammonis and the granular layer of dentate gyrus from analysis since their cells are too densely packed for effective segmentation with our automated pipeline, leaving 41 structures of the hippocampal formation.

Next, the scalable vector graphic (svg) plates corresponding to each structure and section (654 in total) were retrieved through the Allen Brain Atlas Application Programming Interface (RRID:SCR_005984) to use as masks for cropping individual structures from the entire coronal brain images. Using the freeware program Inkscape.org (RRID:SCR_014479), each svg mask was sequentially pasted directly on top of the corresponding Nissl-stained image. The two were then clipped together so that the Nissl image was cropped in the shape of the svg mask and exported. This operation was repeated for every downloaded coronal section and svg plate, so that each of the resulting 654 cropped images represented one cross-section of an individual hippocampal formation structure (Fig. 1).

**Fig. 1.**
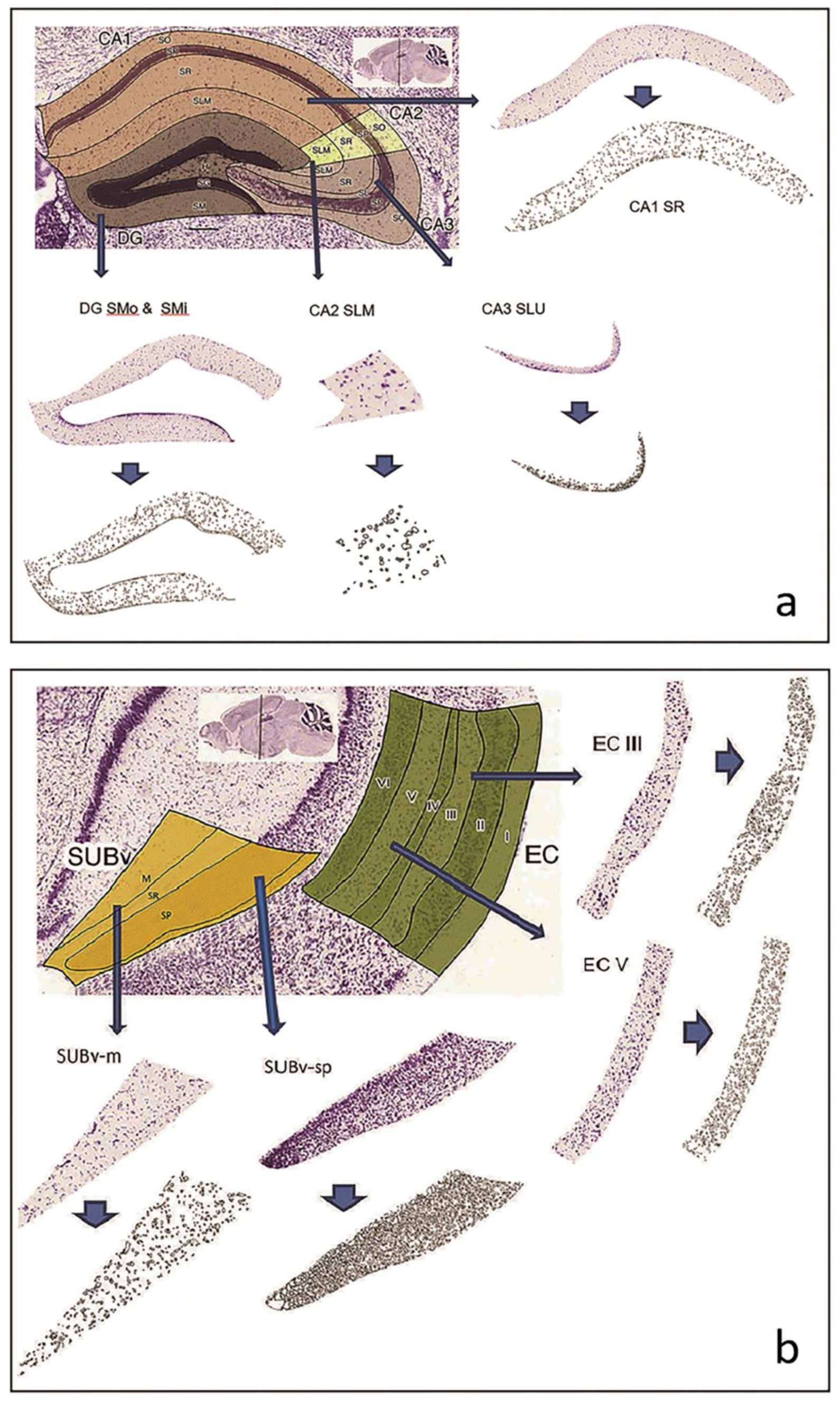
Image source and processing: a. Coronal section of hippocampus areas CA1, CA2, CA3, and Dentate Gyrus (sections illustrated: 74_CA1SR, 74_CA3SLU, 74_CA2SLM and 74_DGmo). b. Coronal section of the entorhinal cortex and the ventral subiculum (sections illustrated: 85_SUBvm and 85_SUBvsp). In both panels, arrows point to the corresponding sample Nissl stained sections as labeled and the smaller arrows point to the segmented images obtained from ImageJ software. Figure was created using ImageJ, Microsoft PowerPoint and Adobe Photoshop

### Image processing

We utilized two distinct image segmentation pipelines in parallel for independent processing and analysis of the acquired images: ImageJ (RRID:SCR_003070) and CellProfiler (RRID:SCR_007358). Both software tools read each of the 654 images and followed a series of steps that resulted in object-by-object segmentation of each image (Fig. 1). The ImageJ pipeline involved increasing contrast by 0.3% (ImageJ-suggested value) and enhancing sharpness (unspecified default parameter), conversion to binary image, and setting the minimum size threshold for object detection to the recommended value of 3 pixels. The CellProfiler pipeline involved using four modules: ‘UnmixColors’ to convert into grey scale and ‘IdentifyPrimaryObjects’ for optimal object identification (Otsu 1979; Sankur 2004) with the default thresholding factor of 3, ‘MeasureObjectSizeShape’ for specifying the objects to be measured, and ‘ExportToSpreadsheet’ for saving the results in the desired format. The image processing scripts, related calibration files, and segmented images are all released in open source (hippocampome.org/phprev/data/ABA_Counts_Database.zip.

Although ImageJ and CellProfiler were remarkably accurate in identifying and segmenting objects from the 2D images as evaluated by visual inspection, the counts could not be taken to reflect cell numbers directly due to four well-known deviations. *First*, several sections exhibited substantial cell clumping, which results in overestimating cell size and underestimating cell count, as each multi-cell clump is generally segmented as a single object. To solve this issue, we applied the corrective watershed algorithm (LaTorre et al. 2013) for separating the clumped cells (Fig. 2a). *Second*, the process of delineating the hippocampal structures within each coronal section splits every border-crossing cell into two, which yields overestimated cell counts and underestimated cell sizes. To alleviate this problem, bordering objects for each section were sorted by area and only the top half were included in the total count for that section, excluding the bottom half (Fig. 2b). *Third*, physical slicing of histological sections inevitably cuts all cells that intersect the surfaces. Since fragments of the dissected cells end up in an adjacent section, cell counts tend to be overestimated and cell sizes underestimated. To address this so-called ‘lost caps’ scenario (Hedreen 1998a), we applied the Abercrombie (Abercrombie 1946) correction (Fig. 2c). *Fourth*, the use of maximum intensity 2D projections through the slice depth risks missing the occluded cells located under those visible from the top view. To adjust the final counts accordingly, we derived a formula by extending Schellart’s theory on count estimates in case of coinciding particles (Hader et al. 2001). Specifically, the tissue can be considered as if divided into “layers” each as thick as the average cell diameter; the number of observed cells (O) then equals the sum of the visible cells in all layers. In the top layer, where there is no occlusion, the number of visible cells equals the real total number of cells (N) divided by the number of layers. In the second layer, the expected fraction of occluded cells corresponds to the areal occupancy of the top layer (the sum of the areas of all cells in the top layer, A, divided by the area of the section (S); thus the number of visible cells equals the real total number of cells (N) divided by the number of layers discounted by that areal occupancy. In the case of two layers, the above reasoning can be summarized in the following equations: 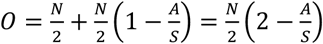, which can be easily solved as: 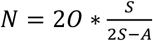. In the case of three layers (Fig. 2d), the expected fraction of occluded cells in the third layer corresponds to the summed areal occupancy of the first and second layers. In this scenario, the formula relating the number of observed cells O to the real number of cells N based on the areal occupancy *A*/*S* is: 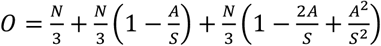, which can be solved as: *N* = 3*O* * *S*^2^/(3*S*^2^ – 3*A* * *S* + *A*^2^).

**Fig. 2.**
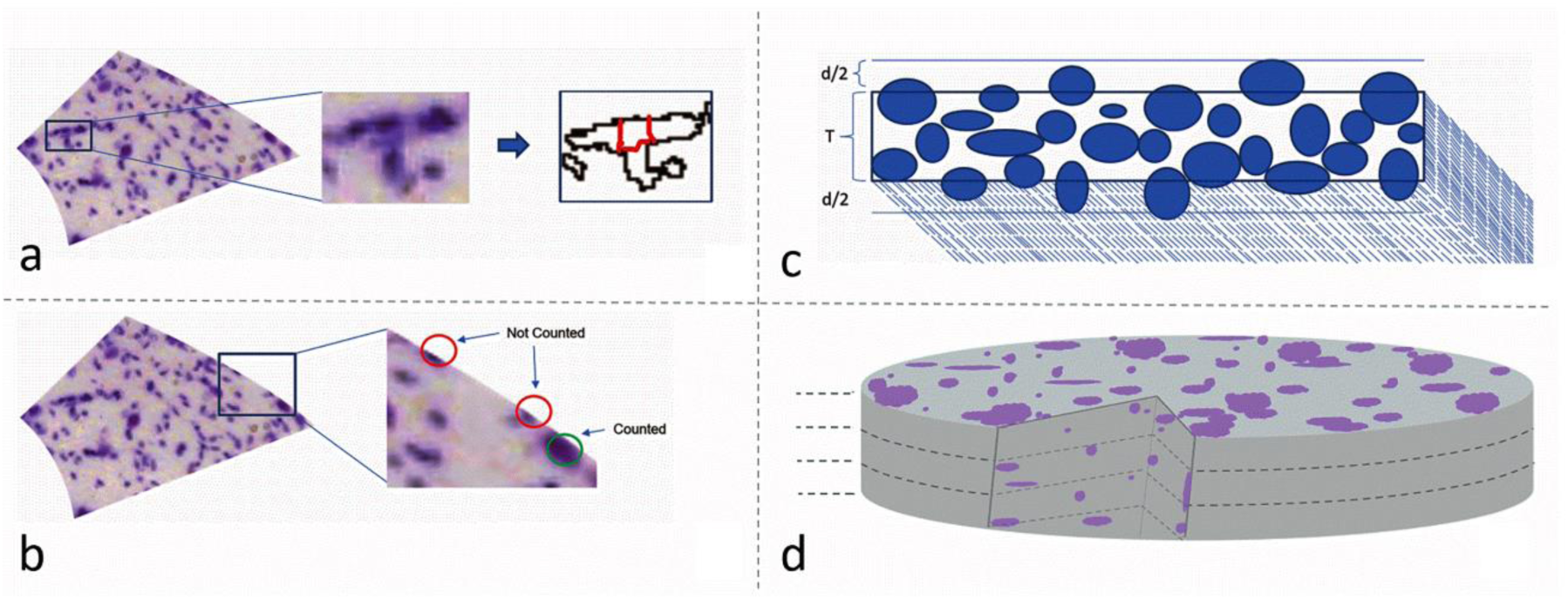
Computing cell counts from segmented object: a. Segmentations were de-clumped using the watershed algorithm (illustrated section: 70_CA2SLM). b. Bordering cells were sorted based on cell area and only upper half was counted for that section while lower half was considered to belong to neighboring areas. c. Cut cells due to sectioning were accounted for using Abercrombie formula: 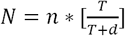, where N is the number of cells after correction, n is the number of all detected objects before correction, T is the section thickness, and d is the mean diameter. d. Section thickness is divided into equal layers where height of each layer equals mean cell diameter; cccluded cells in the depth of the tissue were accounted for using the formula for count estimates aligned particles (see Methods). Figure was created using ImageJ, Microsoft PowerPoint and Adobe Photoshop

### Stereological sampling

In order to compare our computational pipeline to traditional stereological measurements, we counted an unbiased sample of cells in 65 sections (10% of total) using the 2D probe ‘MBF Fractionator’ of the standard commercial software MicroBrightField StereoInvestigator (RRID:SCR_004314). The 65 images were selected randomly by picking 1-2 from each hippocampal structure. For each image, we traced the contour of interest and adjusted the size of the counting frame to allow reliable object identification and accurate marking (this varied for each image depending on the size and distribution of cells). We then specified the size of the Systematic Random Sampling grid based on the section size to ensure a representatively large sample, which is necessary for accurate statistical estimates. Once the grid was placed on the section, we marked each visible object inside the frame or on the inclusion line; after marking all objects in the grid, the process was moved to the next sampling site where the grid was placed. Once all the sampling sites were marked, the process was completed and the probe run list displayed the estimated population for the section. The data files and results were exported and are included in the shared database (hippocampome.org/phprev/data/ABA_Counts_Database.zip).

Shape analysis, bimodality, and spatial distributions. Cell counts were quadrupled since every fourth section from the coronal brain series of Allen’s brain map was Nissl stained and all extracted measurements were converted into metric units multiplying by the reported pixel size of 1.047 µm on each side (Allen Data Production 2011). For each processed image, we extracted two measurements with ImageJ and CellProfiler from every segmented cell: the section area in squared micrometers; and the circularity, defined as 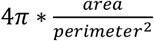, where perimeter is the length of the cell segmentation. While the area quantifies the cell size, circularity characterizes its shape, with a value of 1.0 corresponding to a perfect circle and values closer to 0 indicating increasingly elongated or tortuous shapes.

Analysis was conducted to understand the presence of multimodal distributions in the cell populations based on the size attribute. Kernel density estimation (KDE) plots were generated for each of the 30 parcels of the hippocampus. The KDE plots were fitted with a mixture of two Gaussian distributions, yielding for every parcel a mean and a standard deviation for each of the Gaussians and the relative weight between the two. The proportions of small and large cells per parcel as well as their Gaussian overlap were calculated from these parameters. Hartigan’s Dip test (Maechler 2016) was run on all 30 parcels to test for bimodality/multimodality.

Furthermore, for each processed image we computed three parameters capturing the overall spatial distribution of the cells: the first was volumetric density, defined as the total number of cells in the section divided by the section volume (the product of the mask area by the nominal thickness). The second parameter was the real occupancy (utilized in the occlusion correction described above), defined as the summed area of cells divided by the mask area. The third parameter defines the tiling or clustering tendency of the cells using two-tailed t-test statistics of the nearest neighbor distance (NND) distribution against the null hypothesis (Andrey et al. 2010). Specifically, we randomly distributed within each mask area a number of points identical to that measured in the corresponding section. We then extracted the NND for every point using both the real and random locations. Lastly, we t-tested the real NND distribution against the random NND distribution. A significantly greater NND than random after Bonferroni correction indicates spatial tiling, while a significantly smaller NND than random indicates spatial clustering.

We computed the average and coefficient of variation of the individual cell-level measurements (size and shape) within each section and used the section average in subsequent analyses. We then analyzed all measurements section-by-section rostro-caudally within each hippocampal structure as well as compared the combined sections across structures. The 23 Allen Brain Atlas subdivisions of the medial and lateral regions of the entorhinal cortex were collated into 12 parcels to match the standard nomenclature of Hippocampome.org (Wheeler et al. 2015) as it follows:

- ENTl1 was re-termed LEC I
- ENTl2, ENTl2a, ENTl2b and half the number of cells of ENTl2-3 were combined into LEC II
- ENTl3 and half the number of cells from ENTl2-3 were combined into LEC III
- ENTl4 and half the number of cells from ENTl4-5 were combined into LEC IV
- ENTl5 and half the number of cells from ENTl4-5 were combined into LEC V
- ENTl6a and ENTl6b were combined into LEC VI
- ENTm1 and ENTmv1 were combined into MEC I
- ENTm2, ENTm2a, ENTm2b and ENTmv2 were combined into MEC II
- ENTm3 and ENTmv3 were combined into MEC III
- ENTm4 was re-termed MEC IV
- ENTm5 and half the number of cells from ENTmv5-6 were combined into MEC V
- ENTm6 and half the number of cells from ENTmv5-6 were combined into MEC VI

## RESULTS

Quantitative validation of overall approach. The cell identification process described in the Methods was critically assessed in three distinct ways: by cross-examining the segmentation results from the two independent software frameworks (ImageJ and CellProfiler); by comparing the count results to available data published in the peer-reviewed literature; and by repeating the analysis on a representative subset of the images with traditional unbiased stereology.

Visual inspection of the ImageJ and CellProfiler segmentations against the original images revealed remarkable consistency between the two programs as well as with intuitive evaluation. Specifically, the majority of the objects we would have manually classified with high confidence as cells upon visual inspection were identified as such by both automated pipelines, which also accurately and similarly delineated the body perimeters. When we reviewed a sample of the cells segmented by only one of the two programs and ‘missed’ by the other, most were difficult calls that we could classify as cells only with low confidence. We found no cases of objects identified and segmented by both ImageJ and CellProfiler which we did not consider to be likely cells. On quantitative analysis, the discrepancy in overall cells count between the two programs in the bulk of individual images was within 10%, with an overall difference of less than 103,000 out of a total over 1.5 million. Table 1 reports a summary of these comparisons by anatomical area along with data from published studies (see below). At the level of whole hippocampal formation, the overall count difference between the two software programs was minimal, with an average absolute difference of 7% at the sub-region level. The exact image-by-image counts for both ImageJ and CellProfiler are included in the shared database.

**Table 1.**
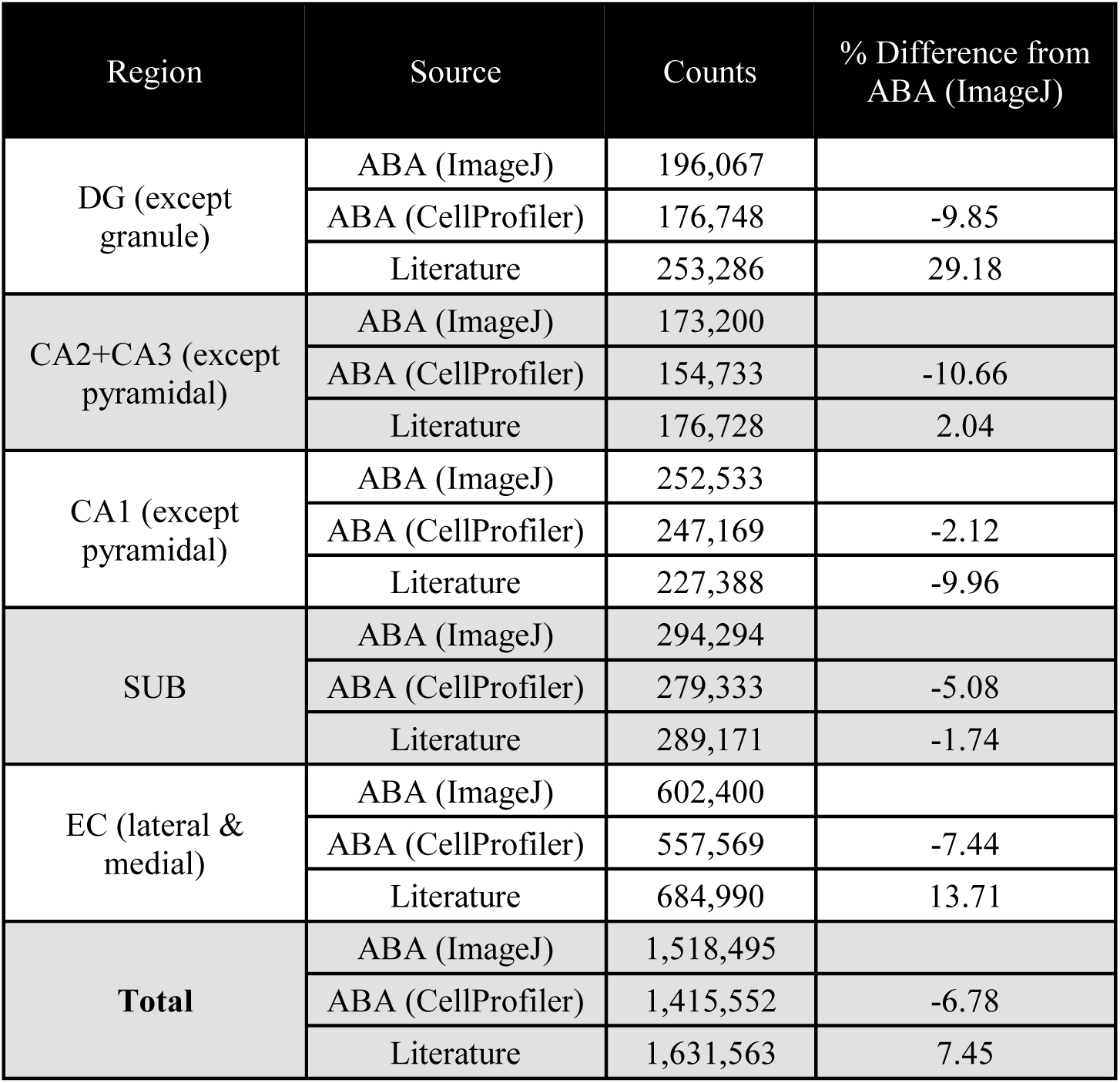
Validation of two independent image segmentation methods with previous studies (Fitting et al. 2009; Mulders et al. 1997; Ramsden et al. 2003; Lister et al. 2006; Sousa et al. 1998; Rasmussen et al. 1996; Kim et al. 2017; Long et al. 1998; Kaae et al. 2012; Murakami et al. 2018; Erö et al. 2018; Bezaire et al. 2016; Andrade et al. 2000; Herculano-Houzel et al. 2013; Grady et al. 2003)

| Region | Source | Counts | % Difference from ABA (ImageJ) |
| --- | --- | --- | --- |
| DG (except granule) | ABA (ImageJ) | 196,067 |  |
|  | ABA (CellProfiler) | 176,748 | -9.85 |
|  | Literature | 253,286 | 29.18 |
| CA2+CA3 (except pyramidal) | ABA (ImageJ) | 173,200 |  |
|  | ABA (CellProfiler) | 154,733 | -10.66 |
|  | Literature | 176,728 | 2.04 |
| CA1 (except pyramidal) | ABA (ImageJ) | 252,533 |  |
|  | ABA (CellProfiler) | 247,169 | -2.12 |
|  | Literature | 227,388 | -9.96 |
| SUB | ABA (ImageJ) | 294,294 |  |
|  | ABA (CellProfiler) | 279,333 | -5.08 |
|  | Literature | 289,171 | -1.74 |
| EC (lateral & medial) | ABA (ImageJ) | 602,400 |  |
|  | ABA (CellProfiler) | 557,569 | -7.44 |
|  | Literature | 684,990 | 13.71 |
| **Total** | ABA (ImageJ) | 1,518,495 |  |
|  | ABA (CellProfiler) | 1,415,552 | -6.78 |
|  | Literature | 1,631,563 | 7.45 |

Comparing the counts obtained with our approach to existing data in earlier scientific studies (mostly using unbiased stereology or nuclear counts) requires a series of assumptions to pool results from a variety of diverse experimental procedures. First, the vast majority of reports on cell counts in the rodent hippocampal formation to date investigated rats rather than mice. In order to relate numerical values between species, we applied the linear scaling parameters reported for cortical structures (Herculano-Houzel et al. 2006): namely, the number of mouse cells in the cerebral cortex (including the hippocampus) equals approximately 34% of the number of rat cells; the number of rat neurons equals approximately 41% of the number of mouse neurons; and the number of rat glial cells equals approximately 29% of the number of mouse glial cells. A summary of the derivation of these scaling rules from the source data is included in the Supplementary Material (Table S1). We utilized the same scaling factors across all sub-regions of the hippocampal formation.

We compiled all other additional assumptions, each specific to the interpretation of individual reports or handling of missing data, in the Supplementary Material as well (Table S2). Lastly, the Supplementary Material (Table S3) also summarizes the step-by-step computations to derive the literature values utilized in Table 1 for each separate sub-region (DG: Table S3a; CA2/3: Table S3b; CA1: Table S3c; subiculum: Table S3d; and entorhinal cortex: Table S3e).

The cell counts obtained with the above-described workflow from both image processing software programs were compared against literature-reported values (Table 1) sourced from 15 distinct studies (Fitting et al. 2009; Mulders et al. 1997; Ramsden et al. 2003; Lister et al. 2006; Sousa et al. 1998; Rasmussen et al. 1996; Kim et al. 2017; Long et al. 1998; Kaae et al. 2012; Murakami et al. 2018; Erö et al. 2018; Bezaire et al. 2016; Andrade et al. 2000; Herculano-Houzel et al. 2013; Grady et al. 2003). The overall differences between our approaches and the average literature data for the whole hippocampal formation overall is remarkably contained (∼8% for ImageJ and ∼15% for CellProfiler). The average absolute difference at the sub-region level (<15%) was considerably smaller (less than half) than the typical variability between experimental studies (Fitting et al. 2009; Mulders et al. 1997; Ramsden et al. 2003; Lister et al. 2006; Sousa et al. 1998; Herculano-Houzel et al. 2013; Grady et al. 2003).

As additional validation of our approach, we subjected a representative subset of the data (∼10% of the images) to unbiased stereology (Fig. 3). This dataset spanned every parcel and ranged from very sparse (<20 cells per image) to considerably dense (>1000 cells per image). Stereological estimates very strongly correlated with our comprehensive counts on the same images (R > 0.99, p < 10^-6^). The overall counts between the two methods differed by less than 5%, and the absolute deviation on an image-by-image case averaged less than 10%. The section-by-section stereological counts are also included in the shared database.

**Fig. 3.**
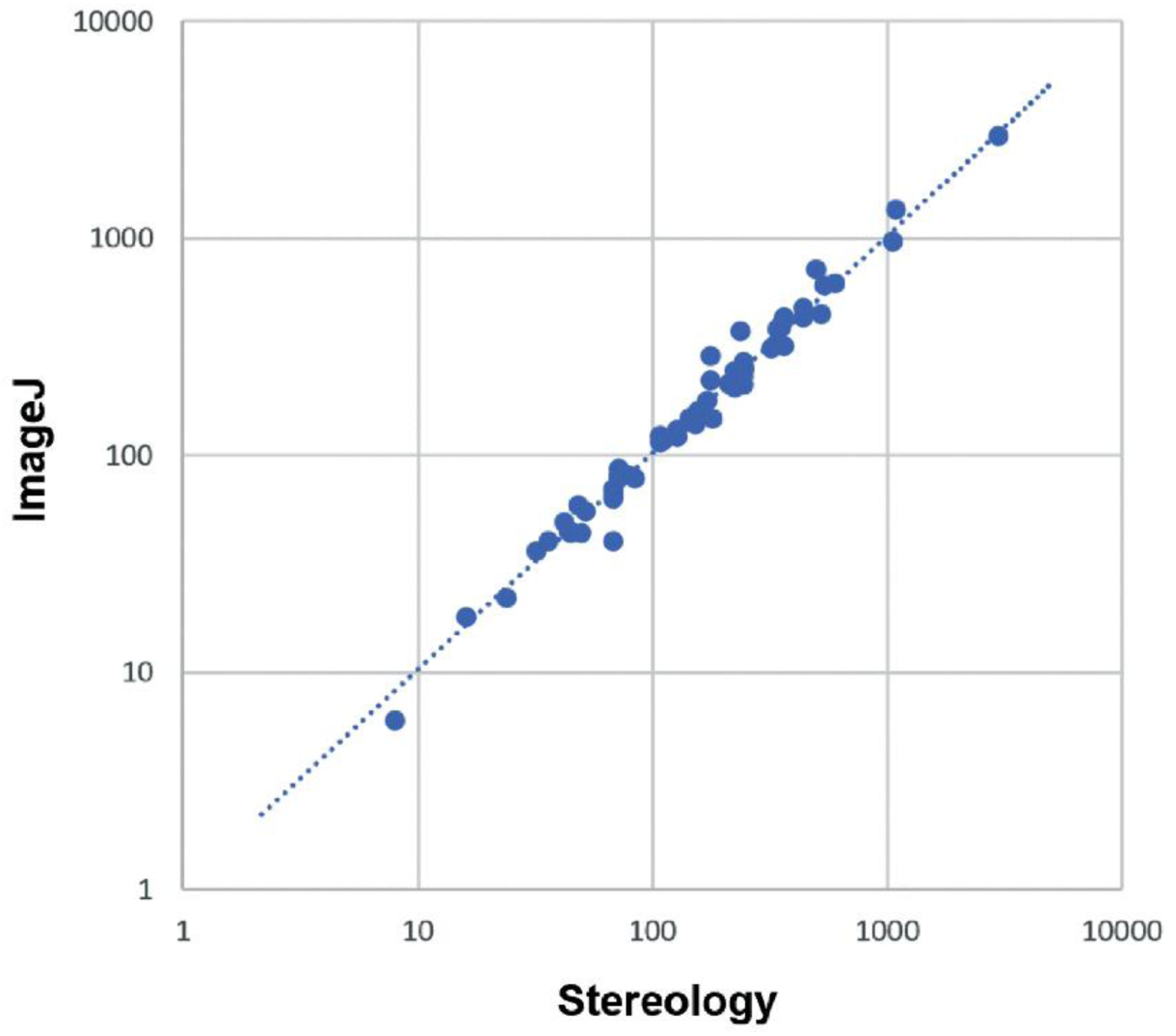
Comparing image processing methods to stereological methods. Figure was created using Microsoft Excel and Adobe Photoshop

Cell numbers, spatial distributions, shape, and size. After validating the analysis pipeline, we collated the measurements of cell count, spatial distribution, shape, and size for each of 30 anatomical parcels (Table 2) to reveal differences and similarities across hippocampal sub-regions and layers.

**Table 2.** Numbers, density, spatial occupancy, size and shape. Area and circularity are reported as averages (values in parenthesis are the coefficients of variation)

| Parcel | Counts | Density (mm<sup>-3</sup>) | Tiling Ratio | Circularity Avg (CV) | Area (μm<sup>2</sup>) Avg (CV) |
| --- | --- | --- | --- | --- | --- |
| DGmo | 155,548 | 75,610 | 3/30 | 0.70 (0.28) | 69.52 (1.16) |
| DGpo | 40,519 | 111,556 | 5/23 | 0.65 (0.28) | 122.08 (0.97) |
| CA3slm | 2,665 | 72,650 | 0/13 | 0.71 (0.25) | 49.65 (0.72) |
| CA3sr | 53,065 | 67,964 | 0/24 | 0.73 (0.26) | 61.65 (0.98) |
| CA3slu | 22,348 | 114,525 | 0/22 | 0.59 (0.36) | 82.95 (1.16) |
| CA3so | 75,530 | 84,499 | 0/24 | 0.68 (0.28) | 67.63 (0.93) |
| CA2slm | 5,656 | 75,103 | 0/15 | 0.74 (0.24) | 50.04 (0.72) |
| CA2sr | 7,136 | 71,148 | 0/16 | 0.72 (0.26) | 64.33 (0.97) |
| CA2so | 6,800 | 81,953 | 0/16 | 0.73 (0.26) | 62.14 (0.95) |
| CA1slm | 94,865 | 80,619 | 2/24 | 0.74 (0.23) | 56.64 (0.77) |
| CA1sr | 82,744 | 58,478 | 0/24 | 0.67 (0.29) | 84.79 (0.93) |
| CA1so | 74,924 | 84,020 | 0/24 | 0.69 (0.26) | 67.08 (0.84) |
| SUBdm | 11,283 | 93,639 | 0/13 | 0.73 (0.26) | 59.11 (0.85) |
| SUBdsr | 10,203 | 73,840 | 1/13 | 0.72 (0.26) | 63.61 (0.84) |
| SUBdsp | 53,216 | 108,550 | 13/13 | 0.66 (0.25) | 133.30 (0.69) |
| SUBvm | 33,832 | 84,815 | 0/11 | 0.74 (0.25) | 51.15 (0.92) |
| SUBvsr | 21,273 | 81,799 | 0/10 | 0.70 (0.27) | 62.96 (0.95) |
| SUBvsp | 164,487 | 119,233 | 14/14 | 0.66 (0.25) | 140.13 (0.70) |
| LEC I | 77,508 | 100,409 | 0/35 | 0.63 (0.38) | 55.59 (1.12) |
| LEC II | 98,750 | 87,268 | 17/53 | 0.65 (0.26) | 154.50 (0.82) |
| LEC III | 62,936 | 77,518 | 9/32 | 0.68 (0.24) | 131.62 (0.77) |
| LEC IV | 28,113 | 63,647 | 7/28 | 0.69 (0.26) | 109.69 (0.75) |
| LEC V | 59,488 | 73,429 | 13/29 | 0.70 (0.23) | 118.16 (0.77) |
| LEC VI | 77,690 | 109,712 | 22/54 | 0.69 (0.23) | 122.04 (0.66) |
| MEC I | 37,217 | 99,948 | 0/19 | 0.69 (0.29) | 47.21 (1.11) |
| MEC II | 66,365 | 126,570 | 13/21 | 0.66 (0.25) | 134.05 (0.90) |
| MEC III | 45,397 | 121,965 | 13/17 | 0.67 (0.25) | 134.78 (0.98) |
| MEC IV | 7,251 | 132,969 | 2/10 | 0.66 (0.28) | 91.63 (0.67) |
| MEC V | 15,585 | 139,513 | 4/14 | 0.67 (0.26) | 104.85 (0.69) |
| MEC VI | 26,101 | 140,819 | 10/13 | 0.67 (0.24) | 111.90 (0.68) |

The number of cells in each parcel varies widely from less than 2700 in stratum lacunosum-molecular of CA3 to more than 160,000 in stratum pyramidale of ventral subiculum. This broad range primarily reflects the known overall volumes of each parcel (for rat hippocampus proper, see e.g. Ropireddy et al. 2012). In addition, we analyzed the results at the individual section-by-section level to investigate any trends in the data within each parcel along the rostro-caudal direction. The relative distribution of cells along the rostro-caudal extent was considerably non-uniform, but tightly corresponded to each relative section area, with a tendency for relatively higher values towards the caudal end for most parcels (Fig. 4). In quantitative terms, the number of cells within each section significantly correlated with that section’s mask area (R=0.9, p<10^-6^). This indicates that volume accounts for more than 80% of the within-parcel rostro-caudal variance in cell numbers. In other words, the volumetric cell density is essentially constant within each parcel throughout the rostro-caudal extent. Between parcels, in contrast, the volumetric cell density ranged broadly from 58,478 mm^-3^ in CA1 stratum radiatum to 140,000 mm^-3^ in layer 6 of medial entorhinal cortex.

**Fig. 4.**
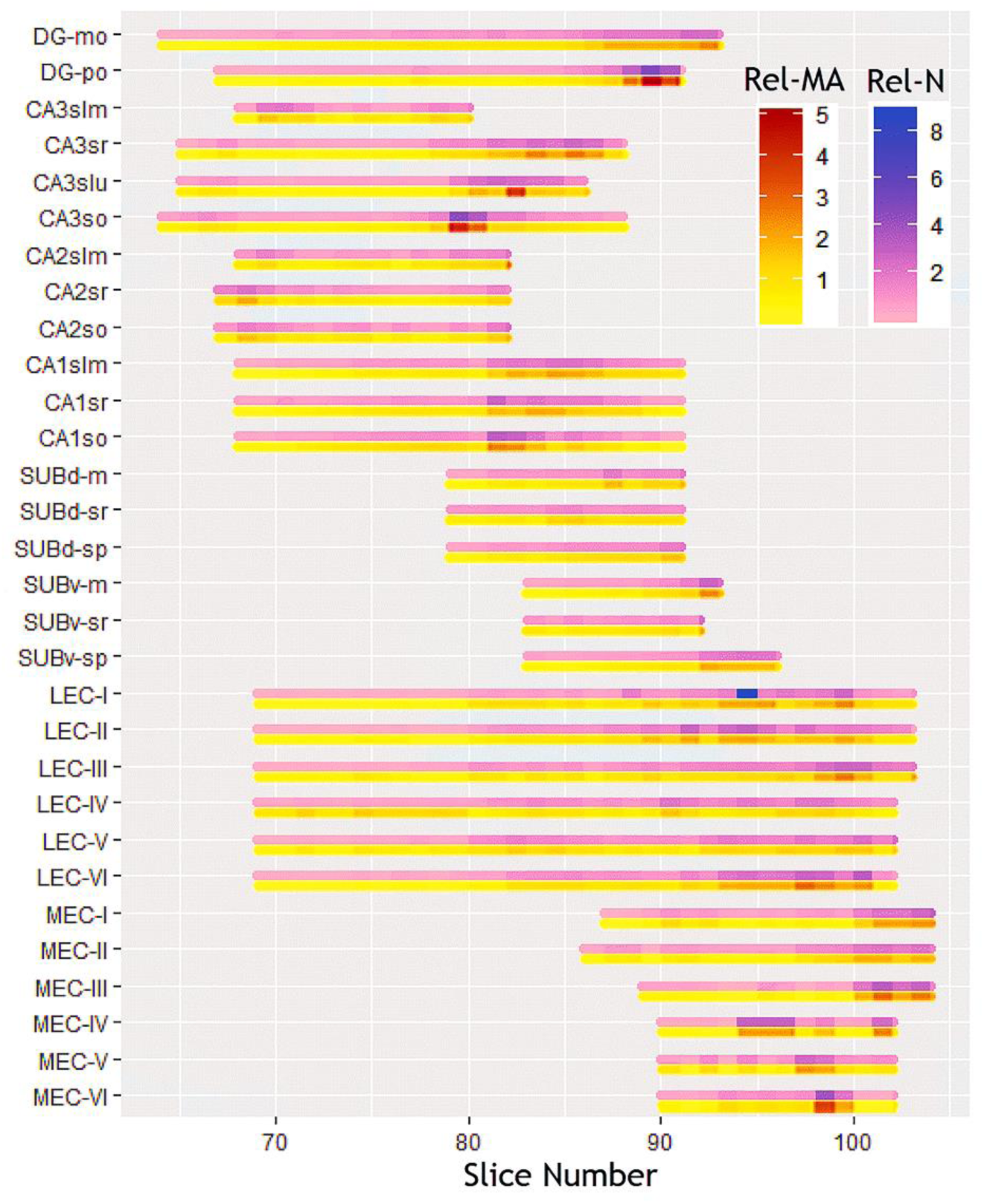
Rostro-caudal gradients of relative cell counts (N) and mask areas (MA). Figure was created using Microsoft Excel, R and Adobe Photoshop

Our statistical analysis of the spatial distributions of cell locations within each parcel was designed to identify two opposite types of deviations from the null hypothesis of random distribution: cell clustering (whereas distances from the nearest neighbors are typically smaller than random) and cell tiling (whereas distances from the nearest neighbors are typically larger than random). None of the 654 sections in our study displayed evidence of clustering (Table 2): 152 (23%) revealed significant tiling (Fig. 5) after Bonferroni correction and 502 (77%) could not be distinguished from random. Only the pyramidal layer in dorsal and ventral subiculum consistently featured cell tiling in all sections. The other subicular layers as well as layer 1 in medial and lateral entorhinal cortex and all CA2 and CA3 parcels had nearly no tiling in any section. The other entorhinal layers and dentate gyrus had an intermediate proportion of tiling sections, ranging from 10% in the dentate molecular layer to 76% in medial entorhinal layer 3, with no observable rostro-caudal trend. Interesting, the proportion of tiling section was significantly correlated to density across the 30 parcels (R=0.72, p<10^-5^), suggesting a greater pressure for optimal space occupancy in cell-denser areas.

**Fig. 5.**
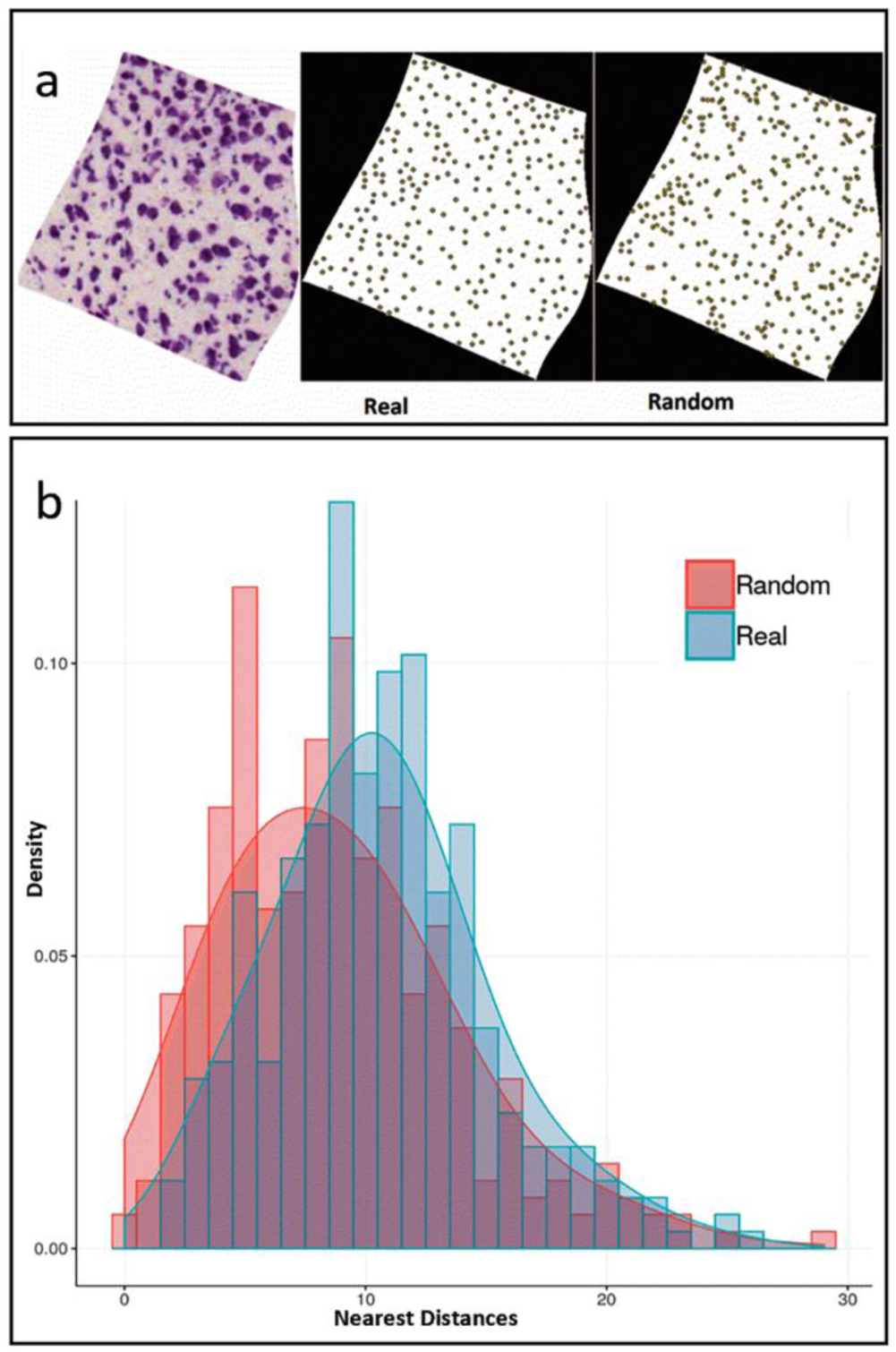
Spatial distributions: a. Nissl stained image (left) for a representative image (79_ENTl4-5), segmentation centroids (center), and randomized points (right); b. Frequency histograms and corresponding probability density functions for real and randomized nearest distances. Figure was created using Microsoft Excel, Microsoft PowerPoint, R and Adobe Photoshop

Cell shape was remarkably uniform throughout the hippocampal formation, with the average circularity only ranging between 0.59 (CA3 lucidum) and 0.74 (stratum lacunosum-moleculare CA1 and CA2, and molecular layer of ventral subiculum) and contained within-parcel coefficients of variations between 0.23 (CA1 lacunosum-moleculare, lateral entorhinal cortex layers V and VI) and 0.38 (lateral entorhinal cortex layer I). Circularity values were very consistent between image analysis pipelines (ImageJ vs CellProfiler) and did not vary along the rostro-caudal axis.

In contrast, cell size varied considerably from parcel to parcel, ranging in section area from less than 50 µm^2^ in medial entorhinal layer 1 to more than 150 µm^2^ in lateral entorhinal layer 2. As a general observation, cells in all layers of areas CA1-3 except CA3 lucidum and CA1 radiatum were twice as small as cells in all layers of the entorhinal cortex with the stark exceptions of both medial and lateral layers 1. The situation was more varied within the dentate gyrus, with cells in the polymorphic layer on average twice as large as in the molecular layer, and in the subiculum, with cells in the pyramidal layers two-and-a-half the size of those in the molecular layer. Cell size was moderately variable within parcel, with coefficient of variations contained between 0.66 and 1.16. No rostro-caudal gradients in cell size were observed in any parcel possibly with the sole exception of lateral entorhinal layer 3, in which rostral cells were marginally larger than caudal ones (similar to micrographs published in (Insausti et al. 1998), though the trend was not statistically significant.

Visual inspection of the Nissl images and corresponding processed segmentations suggested that the cell size distribution within a given parcel and section was typically not regular or uniform, but often skewed, with a substantial number of smaller cells within a more restricted size range and a broader right-tail distributed population of larger cells. Dip test multimodality analysis (Maechler 2016) revealed statistically significant bimodal distributions in a majority of parcels (23/30: Table 3).

**Table 3.** Small cell proportion, misclassified proportion, average areas of small and large cells (values in parenthesis are the coefficients of variation). * denotes significant dip test results

| Parcel | Proportion of small cells % | Misclassified proportion | Small Cells Area ($\mu\text{m}^2$) Avg (CV) | Large Cells Area ($\mu\text{m}^2$) Avg (CV) |
| --- | --- | --- | --- | --- |
| DGmo* | 61.24 | 0.14 | 37.46 (0.33) | 85.81 (0.50) |
| DGpo* | 55.18 | 0.11 | 49.78 (0.54) | 187.17 (0.57) |
| CA3slm | 65.45 | 0.14 | 35.54 (0.30) | 74.46 (0.55) |
| CA3sr* | 73.56 | 0.07 | 37.20 (0.39) | 110.13 (0.51) |
| CA3slu* | 60.00 | 0.13 | 37.15 (0.70) | 139.15 (0.64) |
| CA3so* | 60.70 | 0.11 | 35.78 (0.34) | 105.19 (0.67) |
| CA2slm | 69.00 | 0.10 | 33.96 (0.32) | 76.26 (0.46) |
| CA2sr | 68.18 | 0.11 | 37.27 (0.37) | 95.82 (0.52) |
| CA2so | 58.26 | 0.15 | 32.96 (0.34) | 73.50 (0.48) |
| CA1slm* | 62.00 | 0.14 | 37.77 (0.24) | 72.43 (0.50) |
| CA1sr* | 56.00 | 0.15 | 42.83 (0.42) | 120.97 (0.54) |
| CA1so* | 57.74 | 0.08 | 34.84 (0.31) | 102.76 (0.53) |
| SUBdm* | 59.03 | 0.14 | 35.60 (0.25) | 71.70 (0.46) |
| SUBdsr* | 67.06 | 0.10 | 37.33 (0.34) | 100.78 (0.51) |
| SUBdsp* | 21.61 | 0.16 | 47.19 (0.44) | 153.32 (0.64) |
| SUBvm* | 58.19 | 0.15 | 33.03 (0.32) | 75.07 (0.71) |
| SUBvsr* | 52.00 | 0.15 | 35.47 (0.40) | 100.46 (0.75) |
| SUBvsp* | 19.73 | 0.17 | 54.02 (0.46) | 147.51 (0.58) |
| LEC1* | 62.30 | 0.14 | 28.26 (0.44) | 72.26 (0.54) |
| LEC2* | 20.00 | 0.14 | 46.37 (0.42) | 174.73 (0.63) |
| LEC3* | 23.99 | 0.14 | 42.98 (0.41) | 148.94 (0.59) |
| LEC4* | 39.58 | 0.12 | 41.77 (0.46) | 148.37 (0.51) |
| LEC5* | 41.42 | 0.11 | 47.54 (0.44) | 151.35 (0.46) |
| LEC6* | 75.53 | 0.14 | 92.07 (0.57) | 202.30 (0.48) |
| MEC1* | 67.18 | 0.13 | 27.21 (0.50) | 73.16 (0.68) |
| MEC2* | 62.91 | 0.17 | 75.51 (0.62) | 181.78 (0.43) |
| MEC3 | 76.01 | 0.12 | 91.36 (0.58) | 223.74 (0.47) |
| MEC4 | 19.02 | 0.16 | 33.53 (0.40) | 100.05 (0.55) |
| MEC5 | 53.00 | 0.20 | 63.78 (0.51) | 138.33 (0.46) |
| MEC6* | 76.43 | 0.14 | 77.19 (0.58) | 164.28 (0.42) |

The kernel density estimate (KDE) of all distributions could be well fitted with a Gaussian mixture model (Fig. 6), allowing the determination for every anatomical parcel of the mean and variance of smaller and larger cells, their relative proportion, and the fraction of cells that would assigned to the incorrect group based on a sharp size threshold (“misclassified” area under the cross-over of the two Gaussians). Small cell areas varied considerably between parcels, from less than 30 µm^2^ in medial entorhinal layer 1 to more than 90 µm^2^ in lateral entorhinal layer 6. Similarly, large cell areas also varied over 3-fold between ∼72 µm^2^ in the molecular layer of the dorsal subiculum to ∼224 µm^2^ in layer 3 of the medial entorhinal cortex. The fraction of small cells was balanced overall (average 55%) but ranged widely even across adjacent parcels from more than three-quarters medial entorhinal cortex layer 3 to less than one-fifth in layer 4. The proportion of misclassified cells was relatively modest, ranging from 7% in CA3 radiatum to 20% in medial entorhinal cortex layer 5 (average over all parcels: 13.4%).

**Fig. 6.**
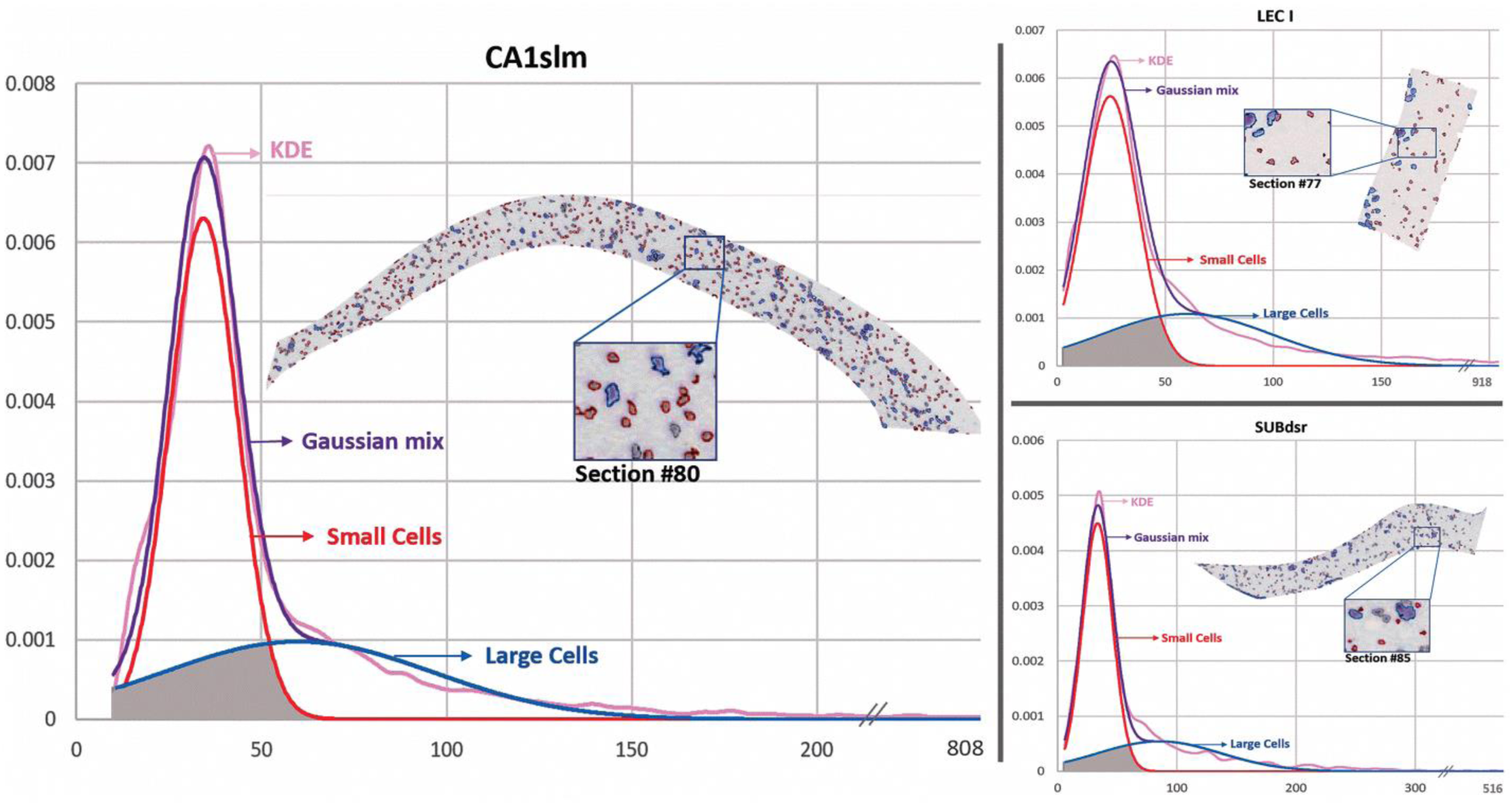
Bimodal cell size distributions of CA1slm (left), LEC I (top right), and SUBdsr (bottom right) with kernel density estimates (KDE) fitted by a Gaussian mixture model. The representative Nissl sections in the insets highlight small and large cells in blue and red, respectively. The gray area under the Gaussian curves represents the proportion of ‘small’ and ‘large’ cells that would be misclassified if selected based on a hard size threshold. Figure was created using Microsoft Excel, Microsoft PowerPoint and Adobe Photoshop

### Data availability

The hippocampome.org/phprev/data/ABA_Counts_Database.zip archive contains all project files and scripts associated with this report, namely: masked segmented images (output files from ImageJ and CellProfiler) for each individual parcel and slice; scripts for corrections and spatial analysis in R; ImageJ scripts for processing, corrections and mask area calculations; CellProfiler calibrations file; and stereology data and results files. The accompanying ‘readme’ document describes each folder and file in detail with instructions on usage. This content is freely available for further research or re-analysis.

## DISCUSSION

The advent of high-throughput histology, accelerating progress in image processing, continuous increase in computing power, and the pervasive accessibility of informatics tools have ushered the era of whole brain cell-by-cell comprehensive analyses into the changing world of neuroscience research. Although there are valid reasons to still favor stereological sampling for certain applications (Schmitz et al., 2014), several recent applications have demonstrated the feasibility of systematically identifying all cell bodies in a mouse brain (Kim et al. 2017; Murakami et al. 2018). Most criticisms of “biased” sampling estimates such as in the Abercrombie formula (Hedreen 1998b) do not apply when considering comprehensive counts across the entire cellular population in a system. Here we show that one of the most popular and widely adopted resources in modern neuroscience, the adult mouse brain Allen Reference Atlas, can be used to obtain cellular-level segmentations throughout entire anatomical formations from the original Nissl stained images. After appropriate corrections to account for sectioning distortions, cell agglomerates, border crossings, and tissue occlusion, we demonstrate that this approach is reproducible across independent image processing pipelines, accurate as matched against unbiased stereology, and reliable in comparison to available published values in the scientific literature.

A key advantage of comprehensive cell segmentation is that, in addition to providing complete counts in each anatomical parcel, it also enables quantitative analysis of cell shape, spatial occupancy, and size. We discovered that, while cross-section circularity (a simple shape measure) is relatively invariant throughout the hippocampal formation, spatial occupancy differs considerably across different parcels, but remains essentially constant within parcels along the entire rostro-caudal extent. Layers 2, 3, 5 and 6 of the medial entorhinal cortex exhibit the highest numerical densities (as also noted in Canto et al. 2008) along with the pyramidal layer of ventral subiculum. This is not surprising since these parcels are especially rich in principal cells, though not in the extreme packing fashion of the dentate granule layer and CA1-3 pyramidal layers, which were excluded for that reason from analysis.

Interestingly, higher occupancy correlated significantly with the fraction of images that demonstrated cell tiling, possibly indicative of pressure towards optimal placement. In contrast, we never observed statistically significant cell clustering. At first, this fact may seem at odd with reports describing cell islands or studies distinguishing island and ocean cells, especially in layer 2 of medial entorhinal cortex. At closer inspections, however, the existing experimental evidence is limited to primates (Goldenberg et al. 1995) or molecularly segregated cell populations (Sun et al. 2015). Seminal studies in rats mention cell clusters, but only in a subset of sections (Insausti et al. 1998). It will be interesting for future studies to ascertain the extent of species specificity in this phenomenon.

Cell size also changed substantially among parcels but was additionally non-uniform within parcels, though again without a discernible rostro-caudal trend. Neuroscientists have maintained a long-standing interest in attempting to distinguish major cell classes on the basis of their somatic sizes. Unfortunately, however, GABAergic cortical interneurons span almost an order of magnitude of cell body areas from approximately 80 µm^2^ to more than 700 µm^2^ (Ascoli et al. 2008), while hippocampal pyramidal neurons fall well within that range with typical somatic areas of the order of 200 µm^2^ (Nakatomi et al. 2002). Thus, it remains untenable to distinguish projection excitatory and local inhibitory neurons in the rodent hippocampal formation solely based on their soma dimension. However, the main classes of glial cells are generally characterized by smaller cell bodies, from microglia in the 20-40 µm^2^ range (Long et al. 1998) to oligodendrocyte and astrocytes in the 40-80 µm^2^ range, respectively (Fitting et al. 2009). Thus, it may be possible at least in principle to separate neurons and glia on the basis of cell size.

The size bimodality analysis presented in this study revealed average areas of 45 µm^2^ and 122 µm^2^ for the “small” and “large” cells, respectively, across the 30 anatomical subdivisions of the hippocampal formation. These values are approximately consistent with the previously reported sizes of glia and neurons; moreover, the overall fraction of small cells (∼55%) is also in line with the expected proportion of glial cells (Herculano-Houzel et al. 2006). Despite these convergent indications, extreme caution needs to be exercised in this interpretation, and the assignment of neurons and glia based on cell size should be deemed tentative at best. With these caveats in mind, it is still worth noting that, based on our segmentation results, layer 1 in medial entorhinal cortex has the smallest cell area across the hippocampal formation. Interestingly, independent experimental evidence suggests that this anatomical parcel is nearly devoid of neurons, whereas the glial density is approximately uniform across layers (Wu et al. 2005; Witter 2011). It is thus tempting to speculate that the number of cells we counted from those sections might correspond mostly or entirely to glia. Glial distribution is also known to vary across the hippocampus: in particular, stratum lacunosum-moleculare displays a higher density of both oligodendrocytes (Vinet et al. 2010) and microglia (Jinno et al. 2007). Intriguingly, this layer also displayed the smallest cell area in all three CA areas.

Direct quantification of distinct cell types (neurons and the three main classes of glia) from Nissl stained images is possible in principle, but requires a higher resolution than currently available in public repositories to enable the examination of physical features of individual cells. At high resolution, specific rules can be applied to identify different cell types such as neurons, astrocytes, and microglia based on size, intensity or texture (Garcia-Cabezas et al. 2016; Rajkowska et al. 2016). Appropriate thresholds for these characteristics can thus be incorporated in existing image processing software tools to efficiently classify objects directly upon segmentation. Similar methodologies can be adapted to quantify any cell type in biological sections as long as appropriate labels are available to mark them. Nonetheless, it is also important to consider that post-mortem handling may alter neuron numbers, stain absorption and selectivity, and other characteristics relevant to the above approach (Gonzalez-Riano et al. 2017). Notwithstanding these caveats, the general method presented here of cell-by-cell segmentation from comprehensive series of stained sections based on readily available image processing software is potentially paradigm-shifting and should be extensible to most of the remaining 700 distinct parcels of the Allen Brain Atlas.

## ACKNOWLEDGEMENTS

We are grateful to Drs. Diek W. Wheeler (from the authors’ lab) and Christopher L. Rees (Harvard University) for critical discussions, and to Dr. Lydia Ng (Allen Institute for Brain Science) for sharing technical details about image scaling and resolution in the Allen Reference Atlas. This work was supported in part by grants R01 NS39600 and U01 MH114829 from the National Institutes of Health.

## REFERENCES

Abercrombie M (1946) Estimation of nuclear population from microtome sections. The Anatomical Record 94:239–247. https://doi:10.1002/ar.1090940210

Allen Data Production (2011) Allen mouse brain atlas technical white paper: in situ hybridization data production. http://help.brain-map.org/download/attachments/2818169/ABADataProductionProcesses.pdf

Andrade JP, Madeira M, Paula-Barbosa M (2000) Sexual dimorphism in the subiculum of the rat hippocampal formation. Brain Research 875:125–137. https://doi:10.1016/s0006-8993(00)02605-6

Andrey P, Kiêu K, Kress C, et al (2010) Statistical Analysis of 3D Images Detects Regular Spatial Distributions of Centromeres and Chromocenters in Animal and Plant Nuclei. PLoS Computational Biology 6. https://doi:10.1371/journal.pcbi.1000853

Ascoli GA, Alonso-Nanclares L, Anderson S, et al (2008) Petilla terminology: Nomenclature of features of GABAergic interneurons of the cerebral cortex. Nature Reviews Neuroscience 9:557–568. https://doi:10.1038/nrn2402

Baldwin SA, Gibson T, Callihan CT, Sullivan PG, Palmer E, Scheff SW (1997) Neuronal Cell Loss in the CA3 Subfield of the Hippocampus Following Cortical Contusion Utilizing the Optical Disector Method for Cell Counting. Journal of Neurotrauma 14:385–398. https://doi:10.1089/neu.1997.14.385

Bahney J, Bartheld CS (2017). The Cellular Composition and Glia-Neuron Ratio in the Spinal Cord of a Human and a Nonhuman Primate: Comparison With Other Species and Brain Regions. The Anatomical Record 301:697–710. https://doi:10.1002/ar.23728

Bayer S, Yackel J, Puri P (1982) Neurons in the rat dentate gyrus granular layer substantially increase during juvenile and adult life. Science 216:890–892. https://doi:10.1126/science.7079742

Bezaire MJ, Raikov I, Burk K, Vyas D, Soltesz I (2016) Interneuronal mechanisms of hippocampal theta oscillations in a full-scale model of the rodent CA1 circuit. ELife 5. https://doi:10.7554/elife.18566

Bezaire MJ, Soltesz I (2013) Quantitative assessment of CA1 local circuits: Knowledge base for interneuron-pyramidal cell connectivity. Hippocampus 23:751–785. https://doi:10.1002/hipo.22141

Bhanu B, Peng J (2000) Adaptive integrated image segmentation and object recognition. IEEE Transactions on Systems, Man and Cybernetics, Part C (Applications and Reviews) 30:427–441. https://doi:10.1109/5326.897070

Boss BD, Peterson GM, Cowan WM (1985) On the number of neurons in the dentate gyrus of the rat. Brain Research 338:144–150. https://doi:10.1016/0006-8993(85)90257-4

Boyce RW, Gundersen HJ (2018) The Automatic Proportionator Estimator Is Highly Efficient for Estimation of Total Number of Sparse Cell Populations. Frontiers in Neuroanatomy 12. https://doi:10.3389/fnana.2018.00019

Bray M, Vokes MS, Carpenter AE (2015) Using CellProfiler for Automatic Identification and Measurement of Biological Objects in Images. Current Protocols in Molecular Biology. https://doi:10.1002/0471142727.mb1417s109

Calhoun ME, Kurth D, Phinney AL, et al (1998) Hippocampal neuron and synaptophysin-positive bouton number in aging C57BL/6 mice. Neurobiology of Aging 19:599–606. https://doi:10.1016/s0197-4580(98)00098-0

Canto CB, Wouterlood FG, Witter MP (2008) What Does the Anatomical Organization of the Entorhinal Cortex Tell Us? Neural Plasticity 2008:1–18. https://doi:10.1155/2008/381243

Erö C, Gewaltig M, Keller D, Markram H (2018) A Cell Atlas for the Mouse Brain. Frontiers in Neuroinformatics 12. https://doi:10.3389/fninf.2018.00084

Fitting S, Booze RM, Hasselrot U, Mactutus CF (2009) Dose-dependent long-term effects of Tat in the rat hippocampal formation: A design-based stereological study. Hippocampus. https://doi:10.1002/hipo.20648

García-Cabezas MÁ, John YJ, Barbas H and Zikopoulos B (2016) Distinction of Neurons, Glia and Endothelial Cells in the Cerebral Cortex: An Algorithm Based on Cytological Features. Frontiers in Neuroanatomy 10. https://doi:10.3389/fnana.2016.00107

Grady MS, Charleston JS, Maris D, Witgen BM, Lifshitz J (2003) Neuronal and Glial Cell Number in the Hippocampus after Experimental Traumatic Brain Injury: Analysis by Stereological Estimation. Journal of Neurotrauma 20:929–941. https://doi:10.1089/089771503770195786

Goldenberg TM, Bakay RA, Ribak CE (1995) Electron microscopy of cell islands in layer II of the primate entorhinal cortex. The Journal of Comparative Neurology 355:51–66. https://doi:10.1002/cne.903550108

Gonzalez-Riano C, Tapia-González S, García A, Muñoz A, Defelipe J, Barbas C (2017) Metabolomics and neuroanatomical evaluation of post-mortem changes in the hippocampus. Brain Structure and Function 222:2831–2853. https://doi:10.1007/s00429-017-1375-5

Häder D (2001) Image analysis: methods and applications. CRC Press, Florida

Hasselmo ME, Stern CE (2015) Current questions on space and time encoding. Hippocampus 25:744–752. https://doi:10.1002/hipo.22454

Hedreen JC (1998a) Lost caps in histological counting methods. The Anatomical Record 250:366–372. https://doi:10.1002/(sici)1097-0185(199803)250:3<366::aid-ar11>3.3.co;2-v

Hedreen JC (1998b) What was wrong with the Abercrombie and empirical cell counting methods? A review. The Anatomical Record, 250:373–380. https://doi:10.1002/(sici)1097-0185(199803)250:3<373::aid-ar12>3.0.co;2-l

Herculano-Houzel S, Lent R (2005) Isotropic Fractionator: A Simple, Rapid Method for the Quantification of Total Cell and Neuron Numbers in the Brain. Journal of Neuroscience 25:2518–2521. https://doi:10.1523/jneurosci.4526-04.2005

Herculano-Houzel S, Mota B, Lent R (2006) Cellular scaling rules for rodent brains. PNAS 103:12138–12143. https://doi:10.1073/pnas.0604911103

Herculano-Houzel S, Ribeiro P, Campos L, Silva AV, Torres LB, Catania KC, Kaas JH (2011) Updated Neuronal Scaling Rules for the Brains of Glires (Rodents/Lagomorphs). Brain, Behavior and Evolution 78:302–314. https://doi:10.1159/000330825

Herculano-Houzel S, Watson C, Paxinos G (2013) Distribution of neurons in functional areas of the mouse cerebral cortex reveals quantitatively different cortical zones. Frontiers in Neuroanatomy 7. https://doi:10.3389/fnana.2013.00035

Hosseini-Sharifabad M, Nyengaard JR (2007) Design-based estimation of neuronal number and individual neuronal volume in the rat hippocampus. Journal of Neuroscience Methods 162:206–214. https://doi:10.1016/j.jneumeth.2007.01.009

Hu T, Xu Q, Lv W, Liu Q (2017) Touching Soma Segmentation Based on the Rayburst Sampling Algorithm. Neuroinformatics 15:383–393. https://doi:10.1007/s12021-017-9336-y

Insausti R, Herrero MT, Witter MP (1998) Entorhinal cortex of the rat: Cytoarchitectonic subdivisions and the origin and distribution of cortical efferents. Hippocampus 7(2):146–183. https://doi:10.1002/(sici)1098-1063(1997)7:23.0.co;2-l

Insausti A, Megías M, Crespo D, et al (1998) Hippocampal volume and neuronal number in Ts65Dn mice: a murine model of down syndrome. Neuroscience Letters 253:175–178. https://doi:10.1016/s0304-3940(98)00641-7

Insel TR, Landis SC, Collins FS (2013) The NIH BRAIN Initiative. Science 340:687–688. https://doi:10.1126/science.1239276

Jinno S, Fleischer F, Eckel S, Schmidt V, Kosaka T (2007) Spatial arrangement of microglia in the mouse hippocampus: A stereological study in comparison with astrocytes. Glia 55(13):1334–1347. https://doi:10.1002/glia.20552

Jones AR, Overly CC, Sunkin SM (2009) The Allen Brain Atlas: 5 years and beyond. Nature Reviews Neuroscience 10:821–828. https://doi:10.1038/nrn2722

Kaae SS, Chen F, Wegener G, Madsen TM, Nyengaard JR (2012) Quantitative hippocampal structural changes following electroconvulsive seizure treatment in a rat model of depression. Synapse 66:667–676. https://doi:10.1002/syn.21553

Kandel ER (2004) The Molecular Biology of Memory Storage: A Dialog Between Genes and Synapses. Bioscience Reports 24:475–522. https://doi:10.1007/s10540-005-2742-7

Kandel ER, Markram H, Matthews PM, Yuste R, Koch C (2013) Neuroscience thinks big (and collaboratively). Nature Reviews Neuroscience 14:659–664. https://doi:10.1038/nrn3578

Kayasandik CB, Labate D (2016) Improved detection of soma location and morphology in fluorescence microscopy images of neurons. Journal of Neuroscience Methods 274:61–70. https://doi:10.1016/j.jneumeth.2016.09.007

Kim Y, Yang GR, Pradhan K, et al (2017) Brain-wide Maps Reveal Stereotyped Cell-Type-Based Cortical Architecture and Subcortical Sexual Dimorphism. Cell 171. https://doi:10.1016/j.cell.2017.09.020

Lamprecht M, Sabatini D, Carpenter A (2007) CellProfiler™: free, versatile software for automated biological image analysis. BioTechniques 42:71–75. https://doi:10.2144/000112257

Latorre A, Alonso-Nanclares L, Muelas S, Peña J, Defelipe J (2013) Segmentation of neuronal nuclei based on clump splitting and a two-step binarization of images. Expert Systems with Applications 40:6521–6530. https://doi:10.1016/j.eswa.2013.06.010

Lau C, Ng L, Thompson C, et al (2008). Exploration and visualization of gene expression with neuroanatomy in the adult mouse brain. BMC Bioinformatics 9:153. https://doi:10.1186/1471-2105-9-153

Lister JP, Tonkiss J, Blatt GJ, Kemper TL, Debassio WA, Galler JR, Rosene DL (2006) Asymmetry of neuron numbers in the hippocampal formation of prenatally malnourished and normally nourished rats: A stereological investigation. Hippocampus 16:946–958. https://doi:10.1002/hipo.20221

Long JM, Kalehua AN, Muth NJ et al (1998) Stereological analysis of astrocyte and microglia in aging mouse hippocampus. Neurobiology of Aging 19:497–503. https://doi:10.1016/s0197-4580(98)00088-8

Luengo-Sanchez S, Bielza C, Benavides-Piccione R, Fernaud-Espinosa I, Defelipe J, Larrañaga P (2015) A univocal definition of the neuronal soma morphology using Gaussian mixture models. Frontiers in Neuroanatomy 9. https://doi:10.3389/fnana.2015.00137

Maechler M (2016) Package ‘diptest’ (Tech.). Retrieved June 15, 2018, from https://cran.r-project.org/web/packages/diptest/diptest.pdf

Malberg JE, Eisch AJ, Nestler EJ, Duman RS (2000) Chronic Antidepressant Treatment Increases Neurogenesis in Adult Rat Hippocampus. The Journal of Neuroscience 20:9104–9110

Meyer HS, Wimmer VC, Oberlaender M, Kock CP, Sakmann B, Helmstaedter M (2010) Number and Laminar Distribution of Neurons in a Thalamocortical Projection Column of Rat Vibrissal Cortex. Cerebral Cortex 20:2277–2286. https://doi:10.1093/cercor/bhq067

Miki T, Satriotomo I, Li H et al (2005) Application of the physical disector to the central nervous system: Estimation of the total number of neurons in subdivisions of the rat hippocampus. Anatomical Science International 80:153–162. https://doi:10.1111/j.1447-073x.2005.00121.x

Moser EI, Moser M, Mcnaughton BL (2017) Spatial representation in the hippocampal formation: A history. Nature Neuroscience 20:1448–1464. https://doi:10.1038/nn.4653

Mulders W, West M, Slomianka L (1997) Neuron numbers in the presubiculum, parasubiculum, and entorhinal area of the rat. The Journal of Comparative Neurology 385:83–94. https://doi:10.1002/(sici)1096-9861(19970818)385:1<83::aid-cne5>3.0.co;2-8

Murakami TC, Mano T, Saikawa S et al (2018) A three-dimensional single-cell-resolution whole-brain atlas using CUBIC-X expansion microscopy and tissue clearing. Nature Neuroscience 21:625–637. https://doi:10.1038/s41593-018-0109-1

Nakatomi H, Kuriu T, Okabe S et al (2002) Regeneration of Hippocampal Pyramidal Neurons after Ischemic Brain Injury by Recruitment of Endogenous Neural Progenitors. Cell 110:429–441. https://doi:10.1016/s0092-8674(02)00862-0

Otsu N (1979) A Threshold Selection Method from Gray-Level Histograms. IEEE Transactions on Systems, Man, and Cybernetics 9:62–66. https://doi:10.1109/tsmc.1979.4310076

Peng H, Roysam B, Ascoli GA (2013) Automated image computing reshapes computational neuroscience. BMC Bioinformatics 14:293. https://doi:10.1186/1471-2105-14-293

Quan T, Zheng T, Yang Z et al (2013) NeuroGPS: Automated localization of neurons for brain circuits using L1 minimization model. Scientific Reports 3. https://doi:10.1038/srep01414

Rajkowska G (2000) Postmortem studies in mood disorders indicate altered numbers of neurons and glial cells. Biological Psychiatry 48:766–777. https://doi:10.1016/s0006-3223(00)00950-1

Rajkowska G, Clarke G, Mahajan G et al (2016) Differential effect of lithium on cell number in the hippocampus and prefrontal cortex in adult mice: a stereological study. Bipolar Disorders 18:41–51. https://doi:10.1111/bdi.12364

Ramsden M, Berchtold NC, Kesslak JP, Cotman CW, Pike CJ (2003) Exercise increases the vulnerability of rat hippocampal neurons to kainate lesion. Brain Research 971:239–244. https://doi:10.1016/s0006-8993(03)02365-5

Rapp PR, Gallagher M (1996) Preserved neuron number in the hippocampus of aged rats with spatial learning deficits. Proceedings of the National Academy of Sciences 93:9926–9930. https://doi:10.1073/pnas.93.18.9926

Rasmussen T, Schliemann T, Sørensen JC, Zimmer J, West MJ (1996) Memory impaired aged rats: No loss of principal hippocampal and subicular neurons. Neurobiology of Aging 17:143–147. https://doi:10.1016/0197-4580(95)02032-2

Ropireddy D, Bachus S, Ascoli G (2012) Non-homogeneous stereological properties of the rat hippocampus from high-resolution 3D serial reconstruction of thin histological sections. Neuroscience 205:91–111. https://doi:10.1016/j.neuroscience.2011.12.055

Sankur B (2004) Survey over image thresholding techniques and quantitative performance evaluation. Journal of Electronic Imaging 13:146. https://doi:10.1117/1.1631315

Schindelin J, Rueden CT, Hiner MC, Eliceiri KW (2015) The ImageJ ecosystem: An open platform for biomedical image analysis. Molecular Reproduction and Development, 82:518–529. https://doi:10.1002/mrd.22489

Schmitz C, Hof P (2005) Design-based stereology in neuroscience. Neuroscience 130:813–831. https://doi:10.1016/j.neuroscience.2004.08.050

Schmitz C, Eastwood BS, Tappan SJ et al (2014). Current automated 3D cell detection methods are not a suitable replacement for manual stereologic cell counting. Frontiers in Neuroanatomy 8. https://doi:10.3389/fnana.2014.00027

Schneider CA, Rasband WS, Eliceiri KW (2012) NIH Image to ImageJ: 25 years of image analysis. Nature Methods 9:671–675. https://doi:10.1038/nmeth.2089

Sherwood CC, Stimpson CD, Raghanti MA et al (2006) Evolution of increased glia-neuron ratios in the human frontal cortex. Proceedings of the National Academy of Sciences 103:13606–13611. https://doi:10.1073/pnas.0605843103

Sousa N, Madeira MD, Paula-Barbosa MM (1998) Effects of corticosterone treatment and rehabilitation on the hippocampal formation of neonatal and adult rats. An unbiased stereological study. Brain Research 794: 199–210. https://doi:10.1016/s0006-8993(98)00218-2

Sun C, Kitamura T, Yamamoto J, et al (2015) Distinct speed dependence of entorhinal island and ocean cells, including respective grid cells. Proceedings of the National Academy of Sciences 112:9466–9471. https://doi:10.1073/pnas.1511668112

Sunkin SM, Ng L, Lau C et al (2012) Allen Brain Atlas: An integrated spatio-temporal portal for exploring the central nervous system. Nucleic Acids Research 41(D1). https://doi:10.1093/nar/gks1042

Tapias V, Greenamyre JT (2014) A Rapid and Sensitive Automated Image-Based Approach for In Vitro and In Vivo Characterization of Cell Morphology and Quantification of Cell Number and Neurite Architecture. Current Protocols in Cytometry 68. https://doi:10.1002/0471142956.cy1233s68

Vinet J, Lemieux P, Tamburri A, Tiesinga P, Scafidi J, Gallo V, Sík A. (2010) Subclasses of oligodendrocytes populate the mouse hippocampus. European Journal of Neuroscience 31(3):425–438. https://doi:10.1111/j.1460-9568.2010.07082.x

West MJ, Slomianka L, Gundersen HJ (1991) Unbiased stereological estimation of the total number of neurons in the subdivisions of the rat hippocampus using the optical fractionator. The Anatomical Record 231:482–497. https://doi:10.1002/ar.1092310411

Wheeler DW, White CM, Rees CL, Komendantov AO, Hamilton DJ, Ascoli GA (2015) Hippocampome.org: A knowledge base of neuron types in the rodent hippocampus. ELife 4. https://doi:10.7554/elife.09960

Witter M (2011) Entorhinal cortex. Scholarpedia, 6:4380.

Wu H, Rassoulpour A, Goodman JH, Scharfman HE, Bertram EH, Schwarcz R (2005) Kynurenate and 7-Chlorokynurenate Formation in Chronically Epileptic Rats. Epilepsia 46:1010–1016. https://https://doi:10.1111/j.1528-1167.2005.67404.x

Zhang D, Liu S, Song Y, Feng D, Peng H, Cai W (2018) Automated 3D Soma Segmentation with Morphological Surface Evolution for Neuron Reconstruction. Neuroinformatics 16:153–166. https://https://doi:10.1007/s12021-017-9353-x

