## Supplementary data for "Cell numbers, distribution, shape, and regional variation throughout the murine hippocampal formation from the adult brain Allen Reference Atlas"

### Brain Structure and Function

Sarojini M. Attili, Marcos F.M. Silva, Thuy-vi Nguyen[#], Giorgio A. Ascoli*

Center for Neural Informatics, Structures, & Plasticity; Krasnow Institute for Advanced Study, George Mason University, Fairfax, VA (USA). [#]Current affiliation: Duke University, Durham, NC (USA).

### Supplementary material: literature analysis.

For rat studies, we used the scaling rules for rodent cortex from physical fractionator work (Herculano-Houzel et al., 2006) to convert counts to mouse (Table S1). Specifically, the following multipliers were used to convert rat data to mouse: 0.34 for pooled neurons and glia; 0.41 for neurons; and 0.29 for glia.

Table S1: Rat to Mouse Scaling (Herculano-Houzel et al., 2006: Cellular scaling rules for rodent brains).

| Species | Total cells ($x10^6$) | % Total cell in cortex | Total neurons ($x10^6$) | % Total neurons in cortex | Total cells in cortex | Total neurons in cortex |
|---|---|---|---|---|---|---|
| Mouse | 108.69 ± 16.25 | 24 | 70.89 ± 10.41 | 20 | $26 \times 10^6$ | $14 \times 10^6$ |
| Rat | 331.65 ± 8.84 | 23 | 200.13 ± 12.17 | 17 | $76 \times 10^6$ | $34 \times 10^6$ |

Table S2: Additional assumptions used to deduce or interpret cell counts values from the literature.

| # | Assumptions |
|---|---|
| 1 | We consider the number of glial cells in the principal layers of CA1, CA2, CA3 and DG as negligible compared to the number of neurons in the same parcels. |
| 2 | We used the combined data from CA2 and CA3 for our comparisons since not enough data exist on these two regions separately for a robust comparison with our numbers. |

| | |
|---|---|
| 3 | We consider Herculano-Houzel’s neuron and non-neuron numbers as corresponding to the reported distribution averages of 400,019 ± 36,262 and 574,736 ± 33,256, respectively. |

Table S3: Multiple source derivation of the average counts used in Table 1 of main text for a. DG; b. CA2/3; c. CA1; d. subiculum; and e. entorhinal cortex.

a. Dentate Gyrus

Neurons

DG-mo

| | | |
|---|---|---|
| Source | Kim et al., 2017 | Ero et al., 2018 |
| Species | Mouse | Mouse |
| Count | 2,817/2 =1,408 (PV+VIP+SST) | 280,099/2 = 140,049 |

DG-po

| Source | Species | Original | Scaled to mouse |
|---|---|---|---|
| Grady et al., 2003 | Rat | 65,420 | 26,822 |
| Fitting et al., 2009 | Rat | 52,495 | 21,523 |
| Mulders et al., 1997 | Rat | 53,000 | 21,730 |
| Ramsden et al., 2003 | Rat | 63200 | 25,912 |
| Lister et al., 2006 | Rat | 49275 | 20,203 |

Glia

Glia

| Source | Species | Cell Type | Original |
|---|---|---|---|
| Long et al., 1998 | Mouse | Microglia | 22,000 (DG-mo + DG-Granule) |
| Ero et al., 2018 | Mouse | Microglia | 84,459/2 = 42,230 |
| Long et al., 1998 | Mouse | Astrocytes | 70,000 (DG-mo + DG-Granule) |
| Ero et al., 2018 | Mouse | Astrocytes | 55,756/2 = 27,878 |
| Ero et al., 2018 | Mouse | Oligodendrocytes | 49,720/2 = 24,860 |

Hilus (DGpo)

| Source | Species | Cell Type | Original | Scaled to mice |
|---|---|---|---|---|
| Grady et al., 2003 | Rat | All glia | 102,000 | 29,580 |
| Kaae et al., | Rat | All glia | 99,600 | 28,884 |

| | | | | | | | | |
|---|---|---|---|---|---|---|---|---|
| | Sousa et al., 1998 | Rat | 45,000 | 18,450 | 2012 (FRL) | | | |
| | Rasmussen et al., 1996 | Rat | 40,000 | 16,400 | Ero et al., 2018 | Mouse | All glia | 65,962/2 = 32,981 |
| Average DG-mo neurons = (1,408 + 140,049)/2 = 70,729 | Kim et al., 2017 | Mouse | 7,289 | | Total Glia: Sum of averages from the above studies | | (32,115+48,939+24,860) + (30,482) = 136,396 | |
| | Ero et al., 2018 | Mouse | 126,772/2 = 63,386 | | Total Glia Dentate Gyrus: ((103,000+94,967)/2) + (30,481) = 129,464 | | | |
| | Average Hilus (DG-po) neurons = 24,635 | | | | | | | |
| Murakami et al., 2018 - all cells (neurons + glia) in DG-mo+DG-po (excluding granule layer) = 549,622/2 = 274,811 | | | | | | | | |
| Average of Murakami and others: ((70,729+24,635+136,396) + 274,811)/2 | | | | | | | | |
| Total DG cells excluding principal layer = 253,286 | | | | | | | | |

| b. CA2/3 | | | | | | | |
|---|---|---|---|---|---|---|---|
| Neurons | | | Glia | | | | |
| Source | Kim et al., | Ero et al., 2018 | Source | Species | Cell Type | Original | Scaled to mouse |
| Species | Mouse | Mouse | Grady et al., 2003 | Rat | All glia | 470,000 | 136,300 |
| Count | 20,060/2=10,030 (PV+VIP+SST) | 127,836 | Ero et al., 2018 | Mouse | All glia | 105,869 | |
| Average of neurons = (10,030 + 127,836)/2 = 68,933 | | | Average of glia = (136,300 + 105,869)/2 = 121,085 | | | | |
| Murakami et al., 2018 - all cells (neurons + glia) in CA2 + CA3 (excluding principal layers) = 326,876/2 = 163,438 | | | | | | | |
| Average of Murakami and others: ((121,085 + 68,933) + (163,438))/2 | | | | | | | |
| Total CA2/3 cells excluding principal layer = 176,728 | | | | | | | |

| c. CA1 | | | | | | | | |
|---|---|---|---|---|---|---|---|---|
| Neurons | | | | Glia | | | | |
| CA1 | | | | CA1 All | | | | |
| Source | Species | Original | Scaled to mouse | Source | Species | Cell Type | Original | Scaled to mouse |
| Bezaire et al., 2016 | Rat | 27,240 | 11,168 (Interneurons) | Grady et al., 2003 | Rat | Astrocytes | 190,000 | 55,100 |
| | | | | Long et al., 1998 | Mouse | Astrocytes | 100,000 | |
| Ero et al., 2018 | Mouse | 345,910/2=172,955 (excluding principal layer neurons) | | Ero et al., 2018 | Mouse | Astrocytes | 59,141/2 = 29,571 | |
| | | | | Grady et al., 2003 | Rat | Oligodendrocytes | 185,000 | 53,650 |
| Kim et al., 2017 | Mouse | 20,572/2= 10,286 (CA1 PV+VIP+SST) | | Ero et al., 2018 | Mouse | Oligodendrocytes | 99049/2 = 49,525 | |
| | | | | Grady et al., 2003 | Rat | Microglia | 95,000 | 27,550 |
| | | | | Long et al., 1998 | Mouse | Microglia | 48,000 | |
| | | | | Ero et al., 2018 | Mouse | Microglia | 118,872/2 = 59,436 | |
| Averaged Neurons of CA1 except principal layer cells = (11,168+10,286+172,955)/3 = 64,803 | | | | Total glia (Sum of the averages of astrocytes, oligodendrocytes and microglia): | | (55,100+100,000+29,571)/3 + (53,650+49,525)/2 + (27,550+48,000+59,436)/3 = 158,140 | | |
| Murakami et al., 2018 - all cells (neurons + glia) in CA1 (excluding principal layer) = 463,665/2 = 231,832 | | | | | | | | |
| Average of Murakami and others: (231,832 + (64,803+ 158,140))/2 | | | | | | | | |
| Total CA1 cells excluding principal layer = 227,388 | | | | | | | | |

| d. Subiculum | | | | | | | | |
|---|---|---|---|---|---|---|---|---|
| Neurons | | | | Glia | | | | |
| Source | Species | Original | Scaled to mouse | Source | Species | Cell Type | Original | Scaled to mouse |
| Mulders et al., 1997 | Rat | 300000 | 123,000 | Fitting et al., 2009 | Rat | Astrocytes | 74,432 | 21,585 |
| Lister et al., 2006 (left side) | Rat | 193725 | 79,427 | Fitting et al., 2009 | Rat | Oligodendrocytes | 159,514 | 46,259 |
| Andrade et al., 2000 (female) | Rat | 300000 | 123,000 | Ero et al., 2018 | Mouse | Astrocytes | 26,212/2 = 13,106 | |

| | | | | | | | |
|---|---|---|---|---|---|---|---|
| Andrade et al., 2000 (male) | Rat | 350000 | 143,500 | Ero et al., 2018 | Mouse | Oligodendrocytes | 65,827/2 = 32,914 |
| Fitting et al., 2009 | Rat | 312252 | 128,023 | Ero et al., 2018 | Mouse | Microglia | 50,869/2 = 25,435 |
| Ero et al., 2018 | Mouse | 435,628/2 = 217,814 | | Astrocyte average + Oligodendrocytes average + microglia | | | ((21,585+13,106)/2) + ((46,259+32,914)/2) + 25,435 = |
| Average of neurons from literature | | | 135,794 | Total glia | | | 82,367 |
| Murakami et al., 2018 - all cells (neurons + glia) in Subiculum = 720,360/2 = 360,180 | | | | | | | |
| Average of Murakami and others: (360,180 + (135,794+82,367))/2 | | | | | | | |
| Total cells in the Subiculum = 289,171 | | | | | | | |

| e. Entorhinal Cortex (Lateral + Medial) | | | |
|---|---|---|---|
| All cells | | | |
| Source | Species | Neurons | Glia |
| Herculano-Houzel et al., 2013 | Mouse | 400,019 | 574,736 |
| Ero et al., 2018 | Mouse | 1,082,511/2 = 541,256 | 626,040/2 = 313,020 |
| Murakami et al., 2018 - all cells (neurons + glia) in Entorhinal Cortex = 910,927/2 = 455,463 | | | |
| Average of Murakami and others: (455,463 + ((400,019+541,256)/2) + ((574,736+313,020)/2))/ 2 | | | |
| Total cells in medial and lateral Entorhinal cortex = 684,990 | | | |
